# Stalk Lodging: A Portable Device for Phenotyping Stalk Bending Strength of Maize and Sorghum

**DOI:** 10.1101/567578

**Authors:** Douglas D. Cook, Witold de la Chapelle, Ting-Che Lin, Shien Yang Lee, Wenhuan Sun, Daniel J Robertson

## Abstract

**Background:** Stalk lodging (breakage of plant stems prior to harvest) is a major problem for both farmers and plant breeders. A limiting factor in addressing this problem is the lack of a reliable method for phenotyping stalk strength. Previous methods of phenotyping stalk strength induce failure patterns different from those observed in natural lodging events. This paper describes a new device for field-based phenotyping of stalk strength called “DARLING” (Device for Assessing Resistance to Lodging IN Grains). The DARLING apparatus consists of a vertical arm which is connected to a horizontal footplate by a hinge. The operator places the device next to a stalk, aligns the stalk with a force sensor, steps on the footplate, and then pushes the vertical arm forward until the stalk breaks. Force and rotation are continuously recorded during the test and these quantities are used to calculate two quantities: stalk flexural stiffness and stalk bending strength.

**Results:** Field testing of DARLING was performed at multiple sites. Validation was based upon three factors. First, the device induces the characteristic “crease” or Brazier buckling failure patterns observed in naturally lodged stalks. Second, in agreement with prior research, flexural stiffness values attained using the DARLING apparatus are strongly correlated with bending strength measurements. Finally, a paired specimen experimental design was used to determine that the flexural data obtained with DARLING is in agreement with laboratory-based flexural testing results of the same specimens. DARLING was also deployed in the field to assess phenotyping throughput (# of stalks phenotyped per hour). Over approximately 5000 tests, the average testing rate was found to be 210 stalks/hour.

**Conclusions:** The DARLING apparatus provides a quantitative assessment of stalk strength in a field setting. It induces the same failure patterns observed in natural lodging events. DARLING can also be used to perform non-destructive flexural tests. This new technology has many applications, including breeding, genetic studies on stalk strength, longitudinal studies of stalk flexural stiffness, and risk assessment of lodging propensity.

## INTRODUCTION

Stalk lodging (breakage of plant stems prior to harvest) reduces grain yields by approximately 5% annually (Duvick, 2005). Crop varieties that are high yielding and tolerant of crowding stress can be especially susceptible to lodging due to increased grain weight and plant height. Increasing the bending strength of plant stems is therefore important to both current food production and to the development of future crop varieties. It is estimated that US farmers lose almost $380 billion per year in lost maize yield due to the problem of stalk lodging (USDA Annual Yield Reports, Duvick, 2005).

Various testing methodologies for predicting stalk lodging resistance have been presented, including crush tests (Thompson, 1964; Zuber and Grogan, 1961), rind penetration tests (Peiffer et al., 2013) and bending tests (Gou et al. 2008; Hu et al., 2013; Jenkins, 1930; Kokubo et al., 1991; Li et al., 2003; Robertson et al., 2014). In addition, several researchers have sought to establish correlations between various chemical or morphological factors of plants and stalk lodging resistance (e.g., measurements of stalk diameter, stalk lignin content, rind thickness, etc.). Unfortunately these prior art methods are typically labor-intensive and time consuming and often require the use of expensive laboratory equipment (e.g., laboratory-based bending and crushing tests), making them unattractive to plant breeders. In addition, the majority of these methods do not produce the same failure types and patterns observed in naturally lodged maize stalks.

The predominant failure pattern of maize stalk lodging is a distinctive crease near the node (Robertson et al., 2015). Prior art methods of measuring stalk strength induce failure patterns that are substantially different from this creasing pattern (see for example, crushing tests: Thompson, 1964; Zuber and Grogan, 1961; rind penetration tests Peiffer et al., 2013; and bending tests Jenkins, 1930, Hu et al., 2013, Gou et al. 2008; Kokubo et al., 1991; Li et al., 2003). The discrepancy between imposed and natural failure patterns limits the utility of previous methods for phenotyping stalk lodging resistance. This paper describes a portable, field-based tool called “DARLING” (Device for Assessing Resistance to Lodging IN Grains) that replicates the failure mode of naturally lodged stalks and provides quantitative measurements of stalk bending strength and stalk flexural stiffness. The authors of the present work are not aware of any other non-proprietary devices for phenotyping stalk bending strength and bending stiffness of large grains in a field setting. The historical lack of such a device has been a significant limitation for breeding efforts focused on reducing stalk lodging.

## DARLING - DEVICE DESCRIPTION

### Overview

The DARLING apparatus consists of three subsystems: frame, sensors, and electronics (i.e., data acquisition and user interface). The frame consists of a vertical arm connected to a footplate via a hinge. The electronics enclosure and the force sensor are attached to the vertical arm, as shown in Figure 1. The method of operation is illustrated in Figure 2. To operate DARLING, the user aligns the stalk with the force sensor, steps on the footplate, presses the “start test” button and pushes the vertical arm forward, causing the stalk to bend (see Figure 1B). Force and rotation are recorded during each test. The recorded data are used to calculate the maximum bending moment applied during the test as well as the flexural rigidity of the stalk. The device components are described in more detail in the following sections.

**Figure 1:**
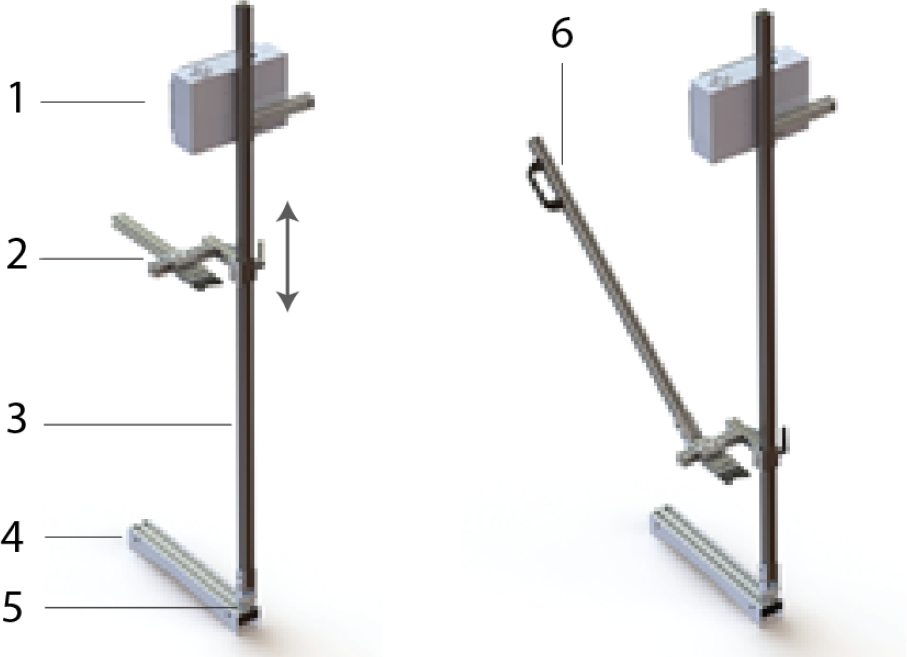
The physical appearance and major components of DARLING. 1 - Electronics and display; 2 - vertically adjustable force gauge; 3 - vertical arm; 4 - footplate; 5 – hinge; 6 - optional handle extension for testing shorter crops or crops that undergo large rotations before failure.

**Figure 2:**
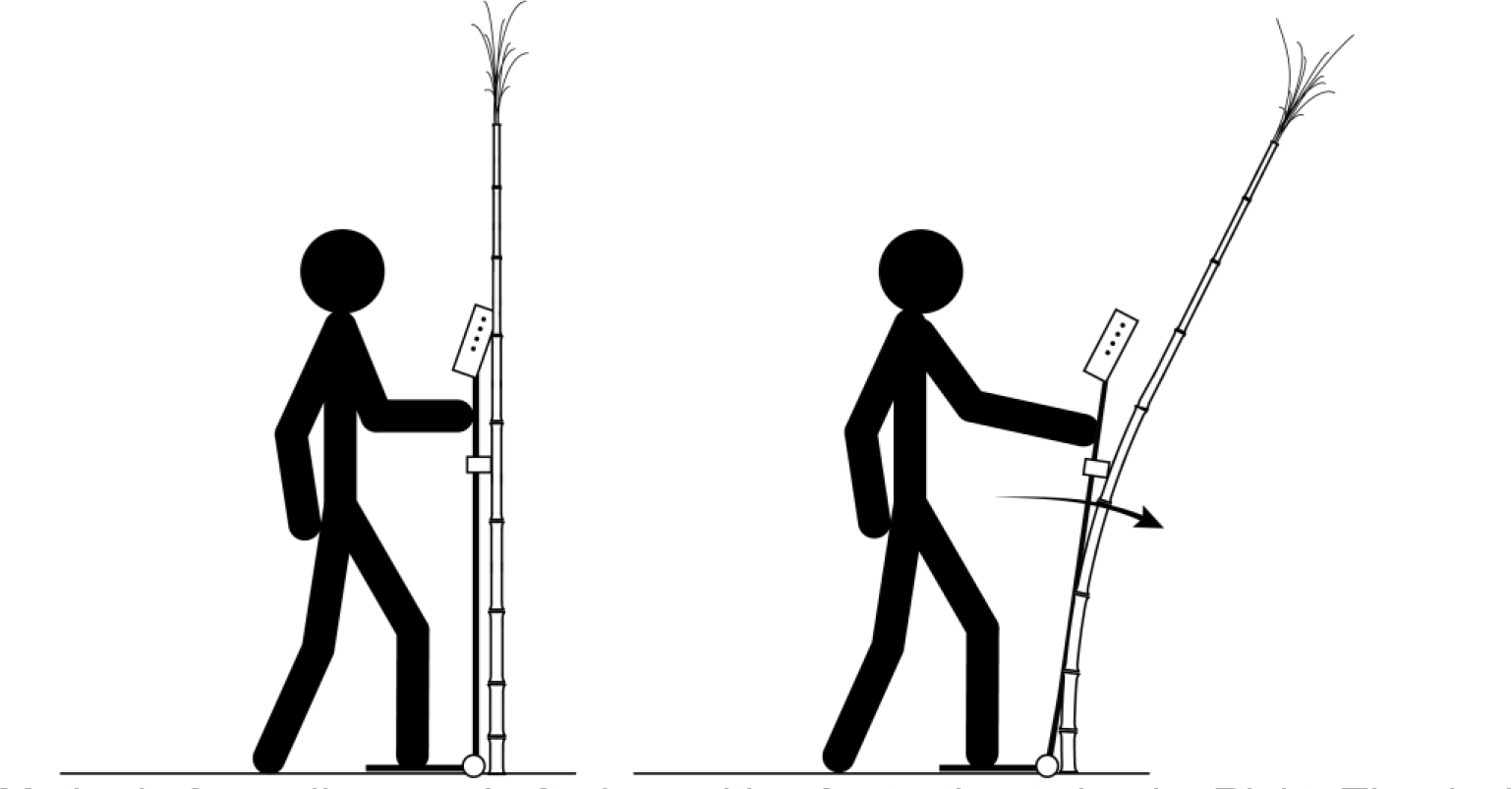
Method of use diagram: Left - In position for testing to begin. Right: The device is pushed forward, inducing bending of the stalk

### Frame

The primary components of the aluminum frame consists of a footplate, a hinge, and a vertical arm. Additional frame components include mounting fixtures for the electronics enclosure and the force gauge. Because crops vary in height, the force gauge mounting bracket is adjustable. Before testing, the operator adjusts the height of the force gauge to match the crop being tested. The operator then reads the height of the force gauge from a scale on the vertical arm, and enters the height into the device’s graphical user interface. An optional arm that provides an ergonomic advantage when testing particularly short crop varieties can also be added to the frame (see Figure 2).

### Graphical User Interface

A graphical user interface is used to control DARLING. The interface consists of a color LCD screen (Adafruit Industries 1933) as well as four selection buttons on the left hand side of the device and four navigation buttons on the top of the device, as shown in Figure 3. The user interface was written using the Python programming language.

**Figure 3:**
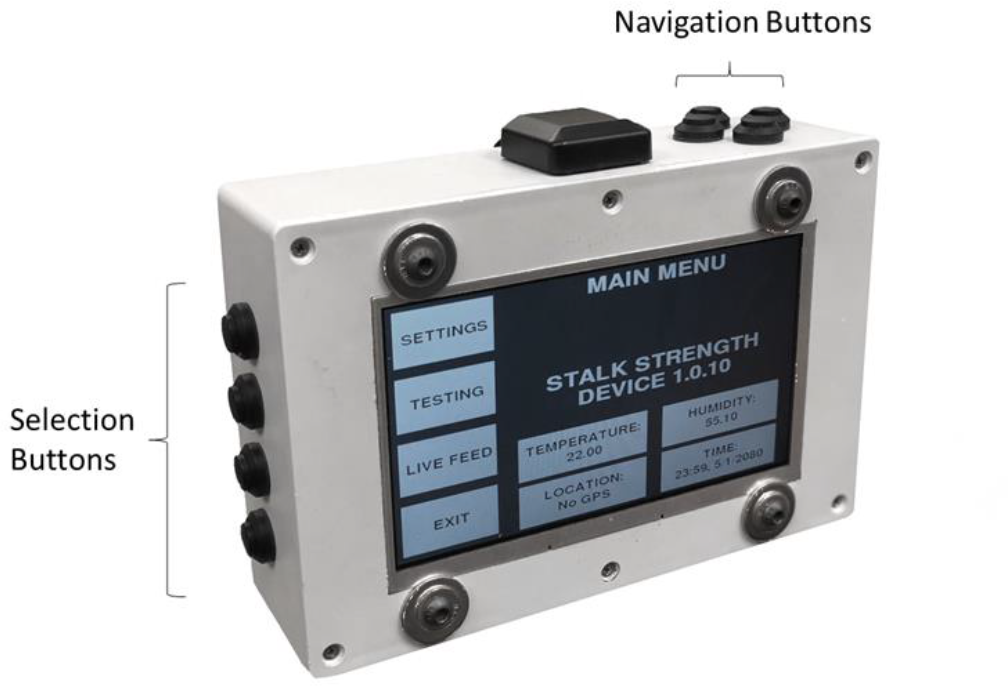
The electronics enclosure and graphical user interface

The user interface provides the following functionality: sensor calibration; real-time sensor verification; a note manager tool; appending of test specific data to each data file (e.g. plot number, plot notes, the height of the force sensor etc.); start/stop test buttons; post-test review of the collected data; software update functionality; data export to a USB drive; and access to the native computer operating system that runs the graphical user interface. Perhaps the most important feature of the user interface is the post-test review screen (Figure 4). This provides the user with a force-deformation plot of each test and allows the user to append additional notes to each test file. At this screen the user also has the option of accepting or rejecting the collected data.

**Figure 4:**
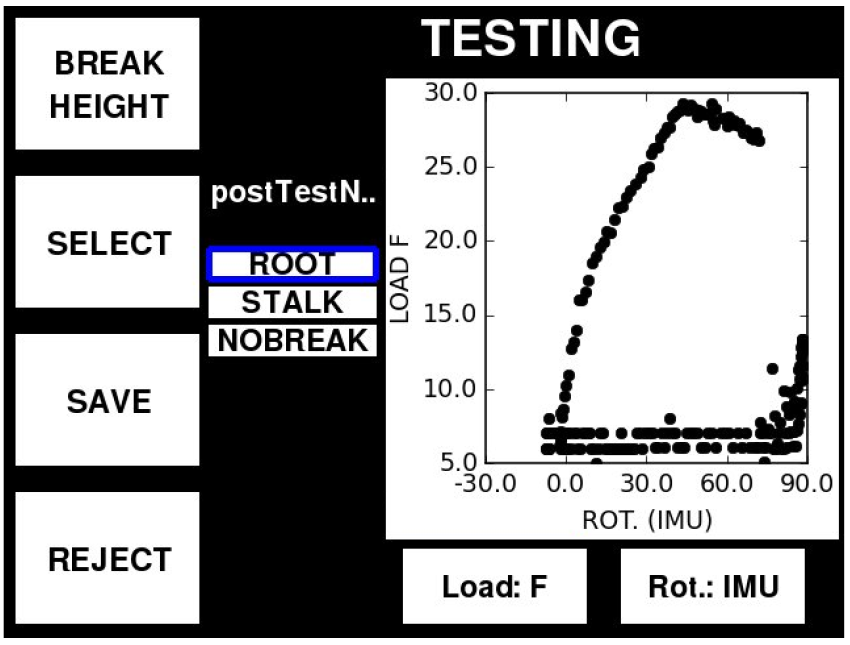
Post-test review screen. The user is able to add post-test notes (center of screen), review the collected data (graph) and save or reject the test data.

### Electronics

The central component of the electronics system is a Raspberry Pi computer (Raspberry Pi RASPBERRYPI3-MODB-1GB) which runs the user interface and records all of the test data. All sensor signals are routed to a microcontroller (Arduino Pro Mini 328, Arduino.cc), which digitizes the signals and forwards the data to the Raspberry Pi computer where it is recorded as shown in Figure 5. DARLING is powered by a lithium Ion battery (PowerCore 20100, Anker) which attaches to the side of the electronics enclosure. The battery can power the device for over 8 hours before requiring a recharge. In addition, the battery is easily accessible for battery exchange in case of a low battery.

**Figure 5:**
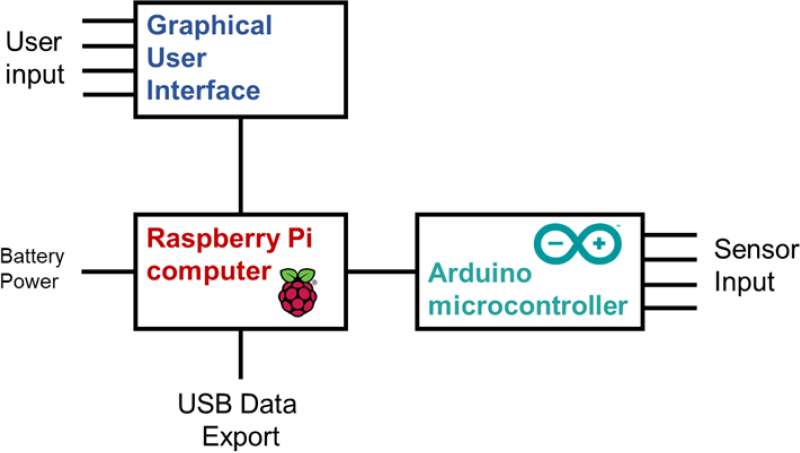
Diagram of major electronic components.

### Data Files

Data is recorded at approximately 30 Hz. Recorded data includes elapsed time, angle of rotation, and force. At the start of each test the ambient temperature, humidity, and GPS coordinates of the device are also recorded. The data from each test is stored in a structured CSV file. The header of the CSV file contains metadata, test attributes, sensor calibration values, and optional data such as field notes. An example data file is provided as supplementary data.

### Post-test calculations

The primary features extracted from the test data are the bending strength (maximum applied bending moment) and the flexural stiffness (i.e., flexural rigidity) of each stalk. The bending strength is computed by calculating the maximum bending moment, *M* at failure:

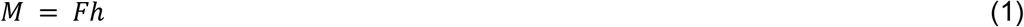

where *F* is the applied force and *h* is the height of the applied force (i.e., the height of the force gauge). The flexural stiffness of the stalk is calculated using the equation for deflection of a cantilever beam and is given by the symbol *EI*:

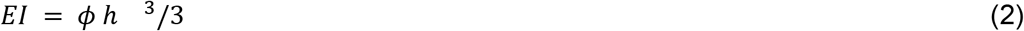

Where *ϕ* is the slope of the linear portion of the force/deformation curve. Deformation is calculated as *h sin(θ)*, where *θ* is the rotational displacement measured by DARLING.

## RESULTS

To date, DARLING has been used to perform several thousand tests on sorghum, and maize at over 15 sites. Researchers from various fields (including, engineering, plant science, genetics, agriculture and high school students) have used DARLING to successfully collect bending strength and bending stiffness data. The results presented below are focused on validation and accuracy of DARLING’s measurements as well as descriptions of the types of data that DARLING produces.

### Failure patterns

DARLING induces failure patterns that are entirely similar to those observed in naturally lodged maize plants. In mature maize plants that have stalk lodged this failure pattern involves creasing of the stem near the node line as described in (Robertson et al., 2015). Mature plants that are diseased or affected by pest damage may exhibit additional failure patterns including splitting or snapping (Robertson et al., 2015). DARLING replicates these failure types and patterns.

Maize plants that have not reached full maturity will often exhibit snapping failures (i.e., the stalk will snap in half) during natural lodging events. This failure type is sometimes referred to as “green snap”. This failure pattern is replicated by DARLING when testing plants that have not reached full maturity.

The last predominant failure pattern observed in natural lodging events is root lodging (uprooting of the plant). This often occurs when plants are rooted in wet or loose soil or when plants exhibit a weak root structure. When testing plants in these soil types DARLING replicates these failure patterns. The user interface of DARLING allows the researcher to either categorize specimens by failure type (root lodged, stalk lodged, splitting, creasing snapping, break height, etc.), or to simply discard failure patterns that are not of interest.

### Validation and Accuracy of Measurements

Several approaches were employed to validate DARLING’s measurements. First data from DARLING was analyzed and compared to typical data patterns observed for structures loaded in bending until failure. We found that data obtained from DARLING exhibits characteristic features common for structures subjected to bending loads to failure, namely: a region of linear force/displacement followed by a departure from linearity, and finally a reduction in load-bearing ability (see Figure 6).

Second, data from DARLING was analyzed to check that it was in agreement with laboratory based protocols for testing maize stalks in bending. In particular, laboratory experiments have demonstrated that stalk flexural stiffness is strongly correlated with stalk bending strength (Robertson et al, 2016). This same correlation is observed in the data collected using DARLING (see Figure 7). In addition, the typical range of failure moments and flexural stiffness values for laboratory based test of maize stalks were compared to the typical range of failure moments and flexural stiffness values acquired using DARLING. We found that both laboratory and DARLING measurements of mature hybrid maize stalks demonstrated failure moments ranging from approximately 3 - 40 Nm.

Third a paired sample experiment was conducted to compare DARLING’s measurements to that of an Instron Universal Testing System (model # 5944, Norwood MA). In particular, DARLING was used to measure the bending stiffness of a number of synthetic samples. Each of these synthetic samples was then tested for bending stiffness using a three-point-bending protocol that utilized the Instron universal testing system. Synthetic samples included wooden dowels, steel rods, and aluminum bars. A total of seven synthetic samples were tested overall. In addition, ten bamboo culms of varying sizes were tested with both the Instron and with DARLING to provide additional points for comparison. DARLING’s flexural stiffness data showed strong agreement with the Instron’s flexural stiffness data. In particular, the coefficient of determination (R^2^) between flexural stiffness data collected with DARLING and flexural stiffness data collected with the Instron was 0.99 for the synthetic samples and 0.98 for the bamboo culms. The flexural rigidity of the samples ranged from 0.3 N-m^2^ to 19 N-m^2^.

**Figure 6:**
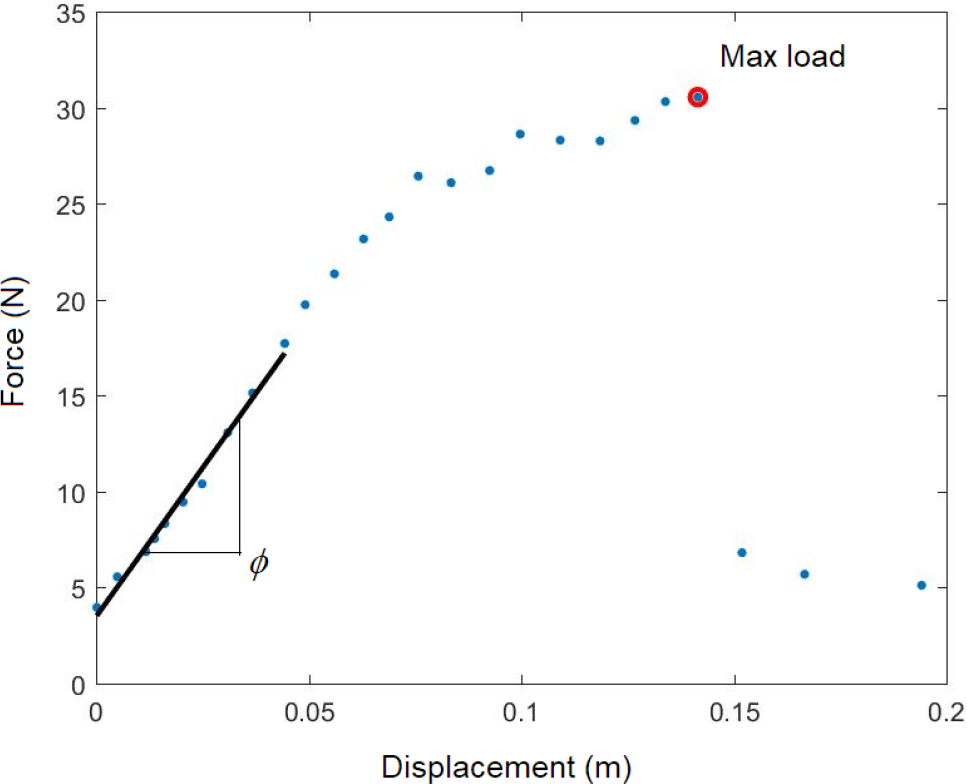
A plot of typical data: applied load plotted vs.displacement. Annotations show the point of maximum load and the regression curve for computing flexural stiffness

**Figure 7:**
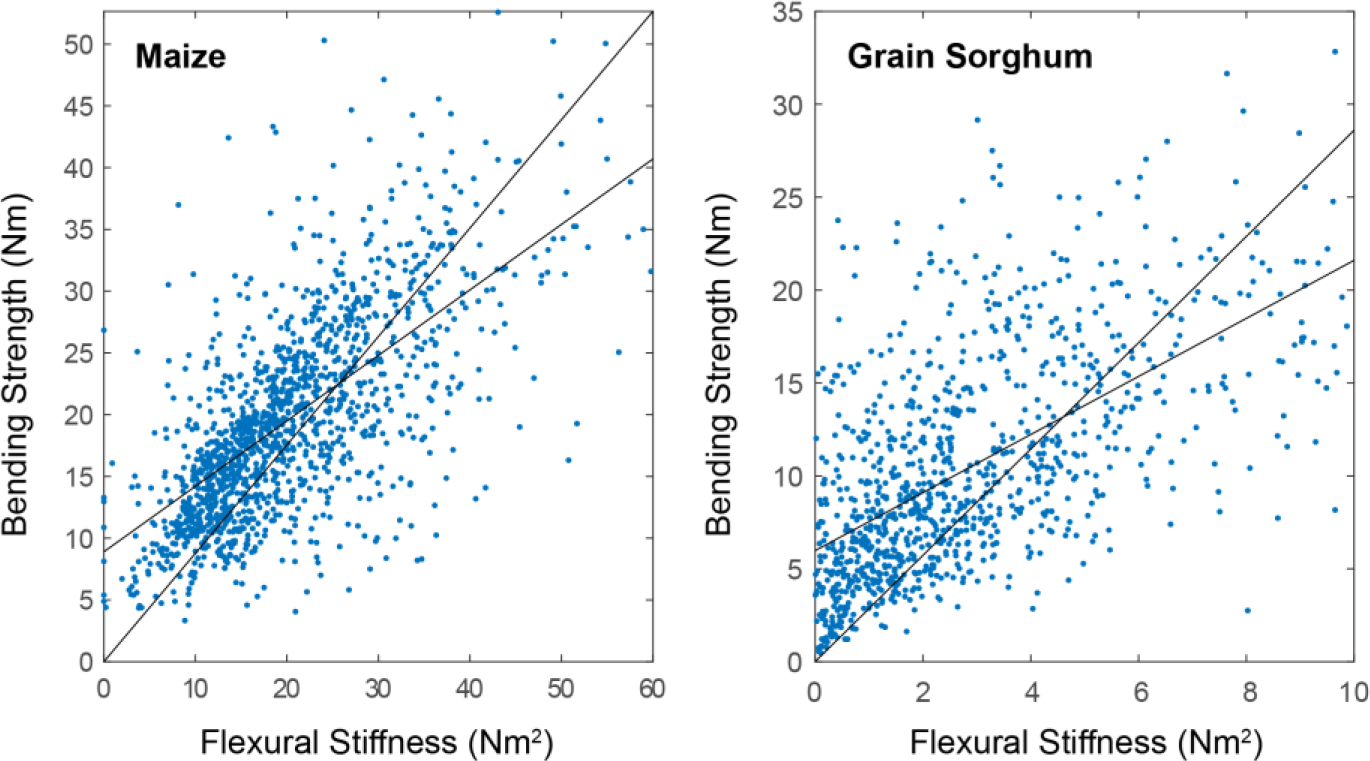
Correlation between stalk flexural stiffness and stalk bending strength was observed in for both maize and sorghum. Regression lines are shown for slope-intercept and slope only regression lines. All regression lines are significant (p-value < 0.001)

### Limitations

DARLING has limitations related to human factors as well as limitations related to technical factors. Limitations related to human factors are presented first, followed by limitations related to technical factors.

Limitations related to human factors, include the labor (and associated cost) required to operate DARLING. Across several thousand tests, and multiple users, the median time required to conduct a single test was 17 seconds, or approximately 210 stalks/hour. The authors are working to implement software changes to DARLING that can automatically detect erroneous test results. This would eliminate the need for the user to inspect the results of each test and make a decision as to the data quality (accept or reject). This improvement should significantly reduce the time required to conduct a test, and simultaneously improve data quality.

The other major human factor limitation is user-to-user variability. Because DARLING is manually articulated, the nature of the applied motion can affect test results. Smooth, gradual rotation of the device leads to cleaner, more reliable data whereas jerky, erratic, and/or rapid motion introduces inertial effects and often leads to poor quality data. Most conscientious users can quickly learn to use DARLING in a consistent manner as test data is immediately displayed following each test.

DARLING also has limitations related to technical factors. These factors include the types of plants that can be tested, and the assumptions used in calculating stalk flexural stiffness and bending strength. With regards to the first technical limitation, DARLING was designed to test large grain plants such as maize and sorghum. In its current form DARLING cannot be used to gather accurate bending strength and flexural stiffness data from small grains such as wheat and barley. The authors are working on a seperate device to measure the bending strength and bending stiffness of small grains.

The second technical limitation of DARLING relates to the equations used to compute stalk flexural stiffness and bending moment. These equations are based on a number of simplifying engineering assumptions. These include the following: (1) when measuring flexural stiffness or bending strength the force is applied perpendicular to the longitudinal stalk axis; and (2) when measuring flexural stiffness (but not when measuring bending strength) the stalk does not rotate in the soil.

Assumption (1), (the force is applied perpendicular to the longitudinal stalk axis), is quite reasonable for small angular deformation values, but becomes less accurate as the stalk is deformed beyond about 20°. In other words, the accuracy of measurements decreases as the amount of rotation increases past 20°. Regarding assumption 1, very accurate results are obtained when testing mature maize stalks in dry soil immediately prior to harvest. In these conditions stalks tend to break when rotated by a fairly small amount. However, stalks that are green (i.e., not fully mature) can bend much further before breaking resulting in decreased accuracy in bending moment measurements. In general, assumption 1 does not affect the accuracy of flexural stiffness measurements. This is because only the first portion of the load displacement curve (the portion occurring prior to 20° of displacement) is used to calculate flexural stiffness.

The validity of assumption (2) (the stalk does not rotate in the soil) depends upon soil conditions and the root structure of the plant being tested. Dry soil with high clay content is often quite rigid, while sandy, wet, or loose soil can allow significant rotation of the stalk and root crown within the soil. Accurate bending strength values can still be obtained in poor soil conditions if the stalks breaks before it has been deflected by 20°. However, flexural stiffness measurements obtained in poor soil tend to underestimate the actual flexural stiffness of the plant stalk.

## CONCLUSIONS

To the best of the authors’ knowledge, DARLING is the first device to induce natural failure patterns and provide reliable measurement of stalk bending strength for corn and sorghum in a field environment. DARLING was developed and thoroughly tested over a period of three years and has proven to be robust and reliable.

DARLING has potential application in breeding studies and for establishing connections between genetics and stalk bending strength. Several such studies are currently underway. The average time to test a single plant with DARLING is approximately 17 seconds/stalk. In an academic or industrial setting the device is capable of testing around 200 stalks/hour, depending on both the individual user and the experimental design.

## REFERENCES

Duvick DN. The Contribution of Breeding to Yield Advances in maize (Zea mays L.). In: Advances in Agronomy [Internet]. Academic Press; 2005 [cited 2018 Jul 12]. p. 83–145. Available from: http://www.sciencedirect.com/science/article/pii/S006521130586002X

Gou L, Zhao M, Huang J-J, Zhang B, Li T, Sun R. Bending mechanical properties of stalk and lodging resistance of maize. Acta Agronomica Sinica. 2008;34(4):653–61.

Hu H, Liu W, Fu Z, Homann L, Technow F, Wang H, et al. QTL mapping of stalk bending strength in a recombinant inbred line maize population. Theoretical and applied genetics. 2013;126(9):2257–66.

Jenkins MT. Experiments on stiffness of stalk. Iowa Agric Exp Stn Annu Rep. 1930;1930:51.

Kokubo, A., Kuraishi S., and Sakurai N., Culm Strength of Barley: Correlation Among Maximum Bending Stress, Cell Wall Dimensions, and Cellulose Content. Plant Physiol. 1989;91(3): 876–882.

Li, K., Yan J, Li J, and Yang X. Genetic architecture of rind penetrometer resistance in two maize recombinant inbred line populations. BMC Plant Biol. 2014;14(1). doi: 10.1186/1471-2229-14-152.

Peiffer JA, Flint-Garcia SA, De Leon N, McMullen MD, Kaeppler SM, Buckler ES. The Genetic Architecture of Maize Stalk Strength. Plos One. 2013 Jun 20;8(6).

Robertson, DJ, Smith S, Gardunia B, and Cook D. An improved method for accurate phenotyping of corn stalk strength. Crop Sci. 2014. 54(5). doi: 10.2135/cropsci2013.11.0794.

Robertson DJ, Julias M, Gardunia BW, Barten T, Cook DD. Corn Stalk Lodging: A Forensic Engineering Approach Provides Insights into Failure Patterns and Mechanisms. Crop Sci. 2015 55(6):2833–41.

Robertson DJ, Lee SY, Julias M, Cook DD. Maize Stalk Lodging: Flexural Stiffness Predicts Strength. Crop Sci. 2016 Aug;56(4):1711–8.

Thompson DL. Stalk Strength of Corn as Measured by Crushing Strength and Rind Thickness. Crop Sci. 1963;3(4):323–9.

Zuber MS, C. O. Grogan. A New Technique for Measuring Stalk Strength in Corn. Crop Sci. 1961;1(5):378–80.

